# Sources and sinks of influenza A virus genomic diversity in swine from 2009 to 2022 in the United States

**DOI:** 10.1101/2025.03.04.641516

**Authors:** Garrett M. Janzen, Blake T. Inderski, Jennifer Chang, Zebulun W. Arendsee, Alicia Janas-Martindale, Mia Kim Torchetti, Amy L. Baker, Tavis K. Anderson

**Affiliations:** Virus and Prion Research Unit, National Animal Disease Center, Agricultural Research Service, U.S. Department of Agriculture, Ames, Iowa, USA; National Veterinary Services Laboratories, National Centers for Animal Health, Animal and Plant Health Inspection Service, U.S. Department of Agriculture, Ames, Iowa, USA

**Keywords:** epidemiology, genetic diversity, influenza A virus, reassortment, swine

## Abstract

Influenza A virus (IAV) in swine in the U.S. is surveilled to monitor genetic evolution to inform intervention efforts and aid pandemic preparedness. We describe data from the U.S. Department of Agriculture National Surveillance Plan for Influenza A Virus in Pigs from 2009 to 2022. Clinical respiratory cases were subtyped followed by sequencing of hemagglutinin (HA) and neuraminidase (NA), and a subset of viruses were whole genome sequenced. Phylogenetic analysis identified geographic and temporal IAV reassortment hotspots. Regions acting as IAV genomic diversity sources or sinks were quantified, and dissemination was qualified and modeled. The dominant IAV clades were H1N2 (1B.2.1), H3N2 (1990.4.a), and H1N1 (H1-1A.3.3.3-c3). Internal genes were classified as triple-reassortant (T) or pandemic 2009 (P), and three genome constellations represented 73.5% of detections across the last two years. In some years, the distribution of IAV diversity was so narrowly distributed that it presented a statistical signal associated with local adaptation. We also demonstrated that the source of most IAV genomic diversity was in Midwest states (IL, MO, IA), and while this was correlated with swine inventory, the emergence and persistence of diversity was tied to swine transport across the U.S. The continued regional detection of unique HA, NA, and genome constellations provides support for targeted interventions to improve animal health and enhance pandemic preparedness.

**Importance:** Variation in the genetic diversity of influenza A virus (IAV) in swine through time and between regions impacts control efforts. This study quantified the genomic diversity of swine IAV collected from 2009-2022 at regional and national levels and modeled sources and sinks of that diversity. Seasonal patterns of IAV transmission were observed, and some locations contributed disproportionately to the emergence of genomic diversity. Minor groups of viruses had the potential to disseminate across the U.S. with animal movement. The identification of these patterns demonstrates the importance of a robust surveillance system to inform vaccine updates that reflect regional patterns of genetic diversity. We show how preemptive interventions in swine IAV diversity hubs could reduce reassortment and the emergence of novel genomic diversity, and how these efforts are likely to reduce the transmission of swine IAV within swine and between swine and humans.

## INTRODUCTION

Influenza A virus (IAV) is a cause of morbidity and mortality in swine resulting in economic loss, and there is a risk for swine IAV to be zoonotically transmitted. In the United States, IAV in swine has been monitored through passive surveillance coordinated by the United States Department of Agriculture (USDA) (1–3). The USDA National Surveillance Plan for Swine Influenza Virus in Pigs seeks to examine the genetic evolution of swine influenza viruses, to inform animal health intervention efforts such as vaccination, and to develop a repository of viruses for research and development (4). The surveillance plan also plays a role in pandemic preparedness as it can detect when and where reassortment between endemic swine IAV and human- and avian-origin IAV (5–7) occurs, a process that not only increases the genetic and antigenic diversity of IAV maintained in pigs, but also increases the probability that a virus with zoonotic potential may emerge (8–10).

Interventions to control IAV in swine are primarily based on biosecurity and vaccines that promote antibodies to the hemagglutinin (HA) (11, 12) protein, with more recent approaches also considering the neuraminidase (NA) (13). Regardless of vaccine platform, matching genetic diversity detected in the field with vaccine components is more likely to result in effective control via reduction in transmission and/or decreased pathology (12, 14). Though a seemingly simple proposition, IAV is a highly diverse RNA virus that is constantly evolving, and in the U.S. there is broad genetic diversity among the HA and NA genes that can vary regionally (3). More than 30 HA genetic clades have been detected (15, 16) among the three endemic swine subtypes (H1N1, H1N2, and H3N2), as well as at least 16 NA genetic clades (16–18). Though there are exceptions, the genetic clades are often antigenically distinct, so constructing an effective polyvalent vaccine is challenging. Consequently, vaccines that are formulated to target diversity within production systems or geographic regions may increase the likelihood of a well-matched and effective vaccine. To achieve this, regular spatial assessment of genetic diversity of swine HA and NA is needed to facilitate laboratory studies that can characterize antibody responses by virus neutralization and hemagglutination inhibition assays to determine when vaccine components should be updated (11, 12, 14).

Significant efforts have been devoted to classifying and describing genetic and antigenic patterns of HA genes in the U.S. (2, 3, 19–23); however, the replication of IAV within, and transmission between, hosts is the result of coordinated function between the HA, NA, and the remaining six “internal” genes (PB2, PB1, PA, NP, M, NS) (24, 25). Internal swine IAV gene constellations in the U.S. are categorized into three groups: triple reassortant origin (TRIG) (26), 2009 H1N1 pandemic (PDM) (27), and related to a live-attenuated influenza virus vaccine (LAIV) in swine (28). Between 2009 and 2016, 70 unique genome constellations were detected among H1 subtypes endemic to U.S. swine (Gao et al. (29)); similarly, Rajao et al. (6) identified at least 44 unique H3 genome constellations; however, in both cases, only 25% of these constellations were maintained across multiple years. Though host immune profile and ecology are likely to play a role, these data suggest that some constellations may have phenotypes associated with improved transmission (30, 31). Data have demonstrated that a human-to-swine spillover H3 HA required pairing with specific internal genes to maintain infection and transmission (5). In addition, subtle differences in gene pairings have been shown to impact virulence (6); for example, reassortment to acquire a PDM-lineage nucleoprotein gene has been shown to improve transmission efficiency (31). Further, reassortment can play a role in antigenic drift and has contributed to the emergence of pandemic viruses in humans (27, 32, 33). Therefore, analyzing the genome diversity of swine IAV may identify patterns that are important in the transmission and evolution of endemic swine IAV, and can inform pandemic preparedness efforts.

In this study, we analyzed genetic sequence data generated through the USDA National Surveillance Plan for Swine Influenza Virus in Pigs from 2009 to 2022. We described spatial and temporal trends in genetic diversity and quantified reassortment patterns across more than 2000 whole genome sequences from samples collected as part of the surveillance plan. We used statistical modeling to detect trends in persistent genome constellations and to elucidate patterns in the generation, dispersal, and IAV diversity across the U.S.

## MATERIALS AND METHODS

### Data acquisition

Genetic sequence data from samples collected as part of the USDA surveillance plan, indicated by a nine-digit alphanumeric barcode beginning with A0 in the virus name, were fetched from octoFLUdb (https://github.com/flu-crew/octofludb). The data were filtered to pure subtype isolates collected from 2009 to 2022 resulting in 10179 viruses with HA and NA gene sequence data (Figure 1). Across this period, a selection of 30 viruses each month were identified for whole genome sequencing (WGS) using a stratified random selection process. Within a given month, the proportion of each HA clade detected was used to determine the number of viruses within that HA clade targeted for sequencing, i.e., if 1A.3.3.3 viruses represented 30% of detections in a month, nine viruses of that clade were randomly selected for WGS. This process resulted in 2632 viruses with WGS data generated between 2009 and 2022. Data from 2023 onwards were not included in this analysis due to a change in the sampling procedure, wherein all samples submitted according to the USDA surveillance plan are now submitted for WGS (16).

**Figure 1.**
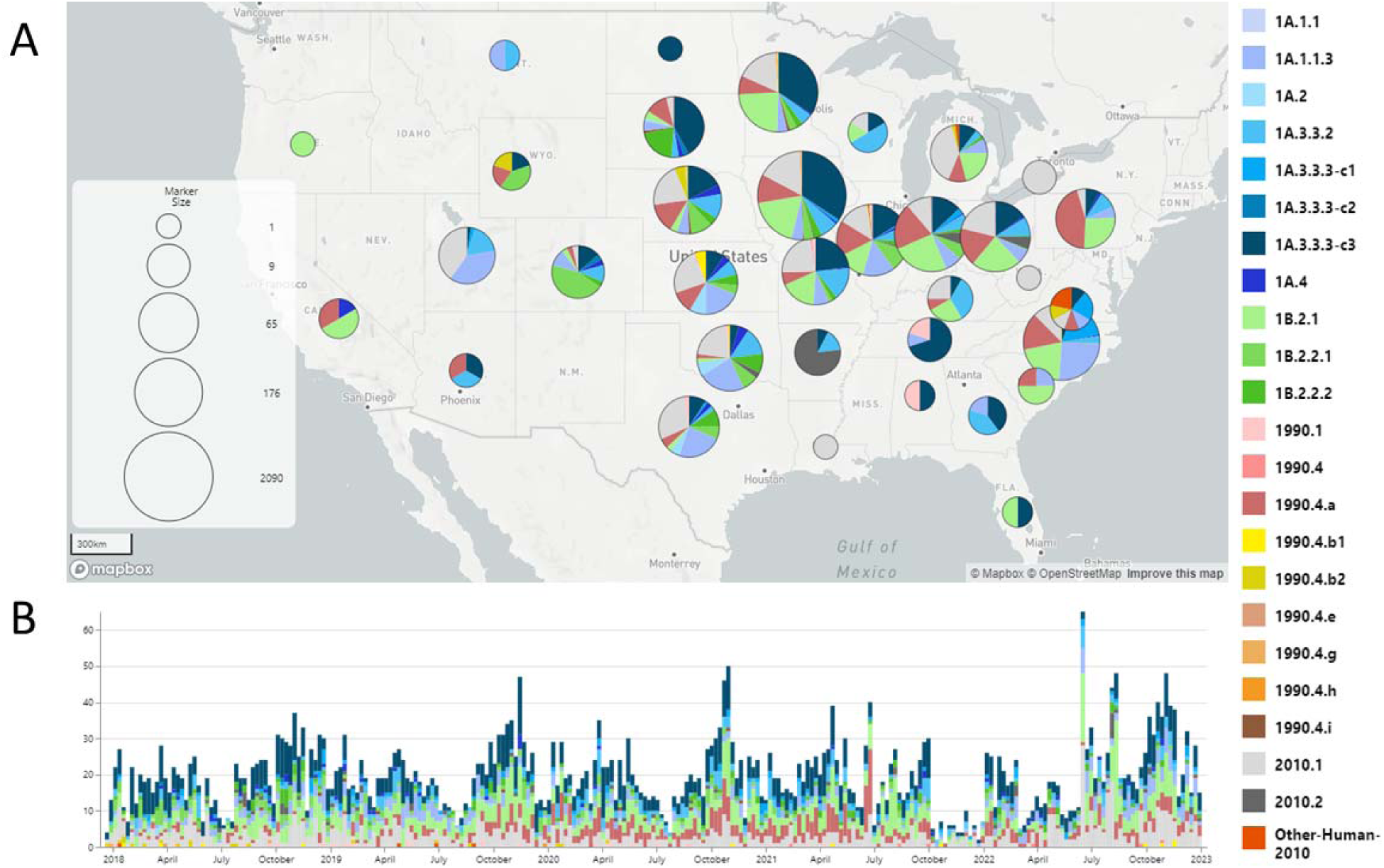
Temporal and spatial visualization of the genetic diversity of influenza A virus (IAV) hemagglutinin gene sequences collected within the United States Department of Agriculture IAV swine surveillance samples between 2017 and 2022. Detections were grouped geographically into states (A) and temporally across the timeline (B).

### Phylogenetic clade classification and quantifying genome constellation diversity

The HA and NA genes were classified to genetic clade with the remaining six genes classified to evolutionary lineage as either triple reassortant (T - TRIG), H1N1 pandemic 2009 (P), or live attenuated influenza virus vaccine (V) (16, 34). Genes were aligned with default settings in MAFFT v7.305b (35), a maximum likelihood phylogenetic tree was inferred using FastTree v2.1.11 (36), and lineage classification was automatically determined using nearest neighbor patristic distances in the octoFLU pipeline (34). The internal gene lineage classification was appended to each virus, generating a genome constellation presented in the order of the six internal genes (PB2, PB1, PA, NP, M, NS). We report trends in HA clade detection frequency and HA/NA clade pairings for: January 2017 to December 2022 to cover unreported data from a prior USDA surveillance assessment (3); and the 2022 calendar year separately to represent more recent trends. Difference plots were generated by determining HA/NA clade pairing counts for single- or two-year periods compared to HA/NA clade pairing counts for the prior single- or two-year counts. Frequency of gene constellation detection from 2009 to 2022 were then identified and summarized using R v3.5.3 and the *ggplot2*, *magrittr*, and *tidyverse* packages.

### Temporal patterns in IAV diversity and persistence

IAV detection frequency was quantified by counting the number of detections per week from 2009-2022. A temporal decomposition (R function *decompose*) was used to quantify broad trends by estimating a moving average, seasonal variation in detection frequency by averaging like-weeks across all years and assigning the remainder of variation to unattributed noise.

All detected genome constellation combinations were plotted on a timeline to identify time periods they were detected. For each unique genome constellation, detection and persistence times were identified by plotting the date on which a constellation was first detected (source event) and its last detection time (sink event). Bars were sorted based on the date of the source event. A six-month window and the beginning and end of the time period were established to demark the period in which we are unable to distinguish between endemic diversity or the emergence or extinction of a novel genome constellation. Consequently, constellations that were detected with first or last detections within these windows at the start and end of the dataset (six months of 2009 and 2022) were not counted as source or sink events, respectively.

### Modeling patterns of interstate spread of IAV genome constellation diversity

To quantify spatial and temporal genetic diversity, we calculated Zeta diversity using the *zetadiv* R package (37) separately for each year between 2009 and 2022. Zeta diversity reflects the average number of states across which each HA clade, NA clade, and genome constellation were dispersed (38). For example, a Zeta score of 5 at order 2 means that five viruses were found in two states, and a Zeta score of 2 at order 5 means that two viruses were found in five states. The diversity index value varies with different combinations of states and decreases as more states are included in a given grouping. Progression through Zeta orders results in the Zeta diversity value decreasing, generating a diversity curve. To ascertain whether units of diversity were dispersed randomly or non-randomly (39), we regressed the ratio of the Zeta decline with either an exponential or power law regression with the Akaike information criterion (AIC) value used to determine which regression best fit the data. A Zeta diversity decline best fit using an exponential regression suggests diversity is driven by random processes, whereas selective processes will produce a pattern that more closely matches the power law regression (38). Sources and sinks of diversity were also quantified by identifying the U.S. state within which a genome constellation was first or last detected. A source-sink score was generated for each state that measured the number of source events less the number of sink events. Each state’s source, sink, and source-sink scores were plotted along with total hog inventory.

To assess whether a genome constellation would disseminate from its initial detection location, we compiled a listing of when and in which U.S. state each genome constellation was detected. Genome constellations that terminated and were never detected again had one final state, “XX,” appended to the end of their state sequence. We used these temporally-ordered U.S. state genome constellation detections to build a discrete-time Markov chain model using the *markovchain* package in R. The transition matrix of this Markov chain provides the probability of where the next detection of a given genome constellation combination will be found given the state in which it was previously detected, as well as the probability of the genome constellation failing to be detected in another state across a 180-day window (sink event).

A network representation plot of the transition matrix was created to visually trace probable routes of interstate transmission between ten states with the highest swine populations, plotting edges with a probability ≥0.15. To assess the robustness of the model, four Markov chain models were trained on a randomly drawn 75% of the data and tested on the remaining 25%. This model’s transition matrix was used to predict the state of final detection, the final novel state transmitted to, and the probability of sink events given different scenarios. Percentage of correct predictions were used to estimate accuracy.

## RESULTS

### HA and NA diversity within USDA IAV swine surveillance samples

From 2009 to 2022, there were 10060 viruses isolated with HA and NA gene sequence data shared publicly through the USDA National Surveillance Plan for Influenza A Virus in Pigs. The subtypes detected were H1N1 (35.6%), H1N2 (35.1%), and H3N2 (28.7%), along with 26 H3N1 and 1 H4N6 virus (Table 1). The most commonly detected HA and NA gene pairings were H1-1B.2.1/N2-1998B (12.2%), H3-1990.4.a/N2-2002B (9.87%), and H1-1A.3.3.3-c3/N1-C.3.2 (9.16%), representing 31.2% of all combinations. Within 2022 alone, the most frequently detected HA genetic clades were H3-2010.1, H1-1A.3.3.3-c3, and H1-1B.2.1; and these three HA genes represented 59.6% of all detections. An additional 13 HA clades represented the remaining 40.4% of detections. The predominant HA/NA gene pairings in 2022 were H3-2010.1/N2-2002B (20.7%), H1-1B.2.1/N2-1998B (17.7%), and H1-1A.3.3.3/N1-C.3.2 (16.5%) (Figure 2). In the past four years, the H1-1A.1.1.3 clade had reassorted: initially the 1A.1.1.3 HA gene was paired with an N2 gene (N2.1998B or N2.2002B), but more recently, this HA gene was most frequently detected paired with an N1 gene (N1-C.3.2 and N1-C.2.1). From 2021 to 2022, the N1-C.3.1 was mostly replaced by N1-C.3.2 in pairings with H1-1A.3.3.3-c3, the H3-1990.4.a/N2-2002B had reduced detection frequency, and the H3 2010.1/N2-2002B pairing was detected more frequently.

**Table 1.**
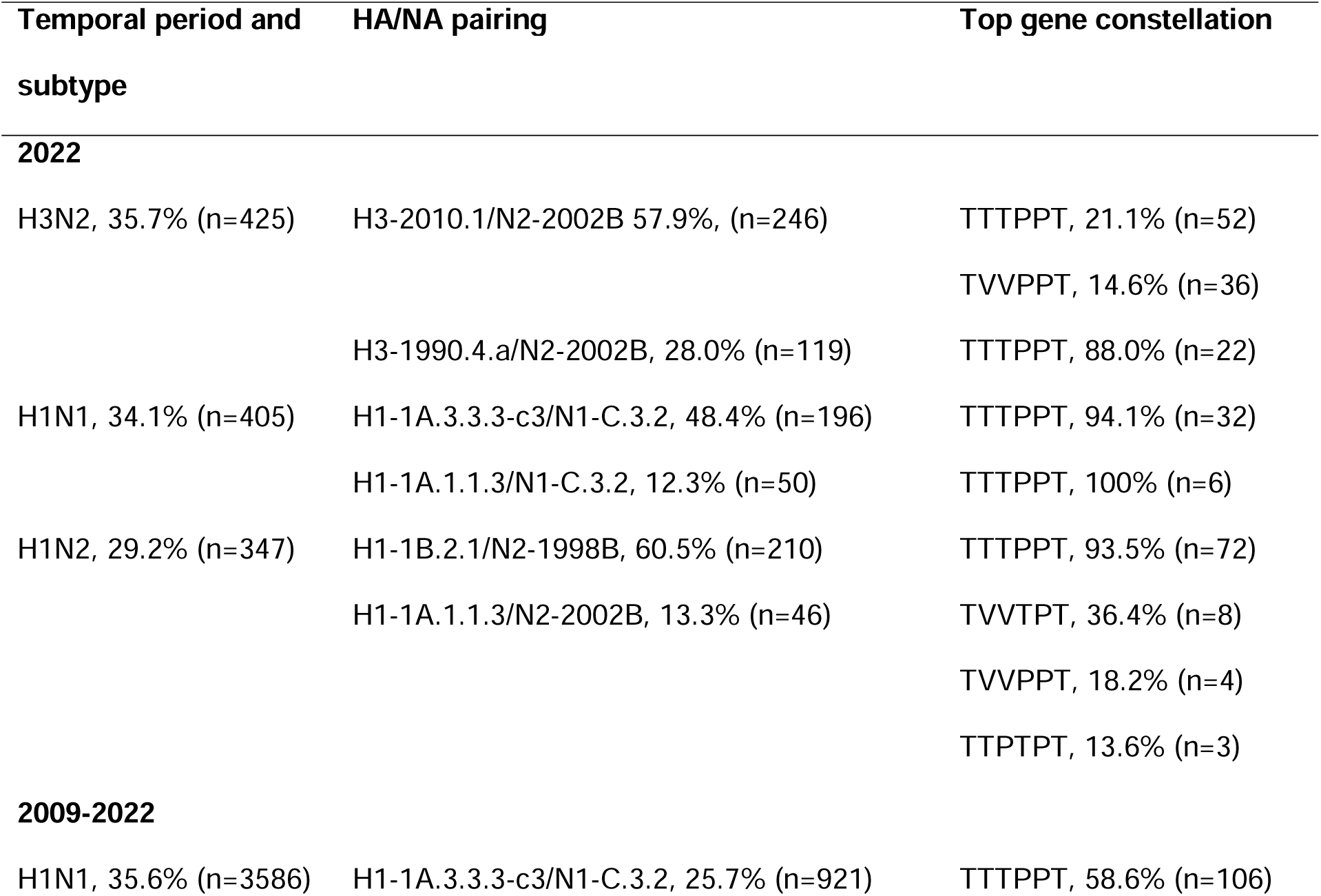
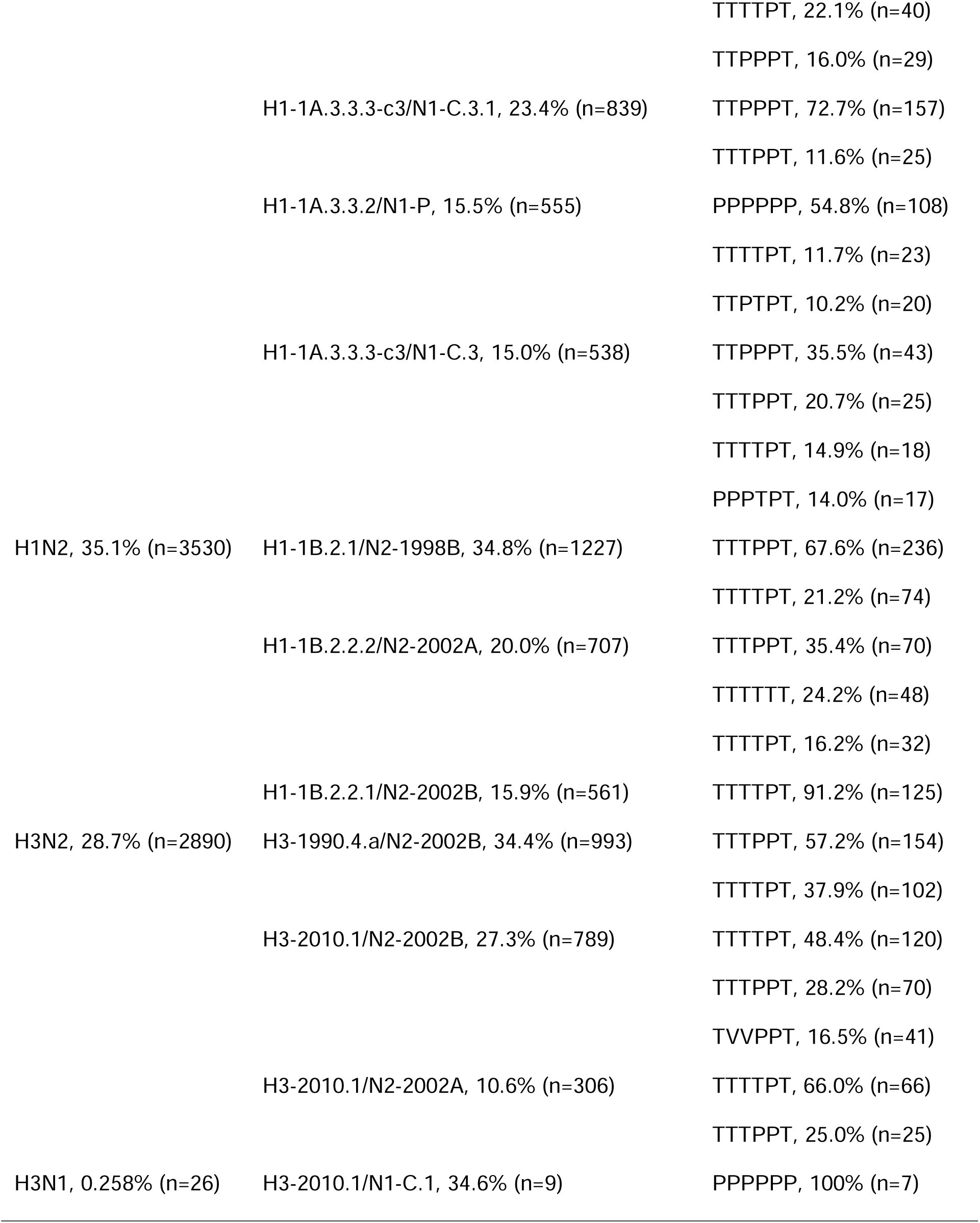
Influenza A virus (IAV) in swine subtype detection frequency, hemagglutinin (HA) and neuraminidase (NA) detection frequency, and associated internal gene constellation detection frequency. The percentages for the HA/NA were calculated within the subtype, and the percentages for the gene constellations were calculated based on the number of whole genomes available (n=359 in 2022; n=2632 between 2009 and 2022). The gene constellation represents the concatenated evolutionary lineage (triple-reassortant – T; H1N1 pandemic – P; LAIV – V) of IAV genes in order: polymerase basic 2 (PB2), polymerase basic 1 (PB1), polymerase acidic (PA), nucleoprotein (NP), matrix (M), and non-structural (NS). Detection frequency was presented for the most recent complete year and for the entire dataset between 2009 and 2022.

**Figure 2.**
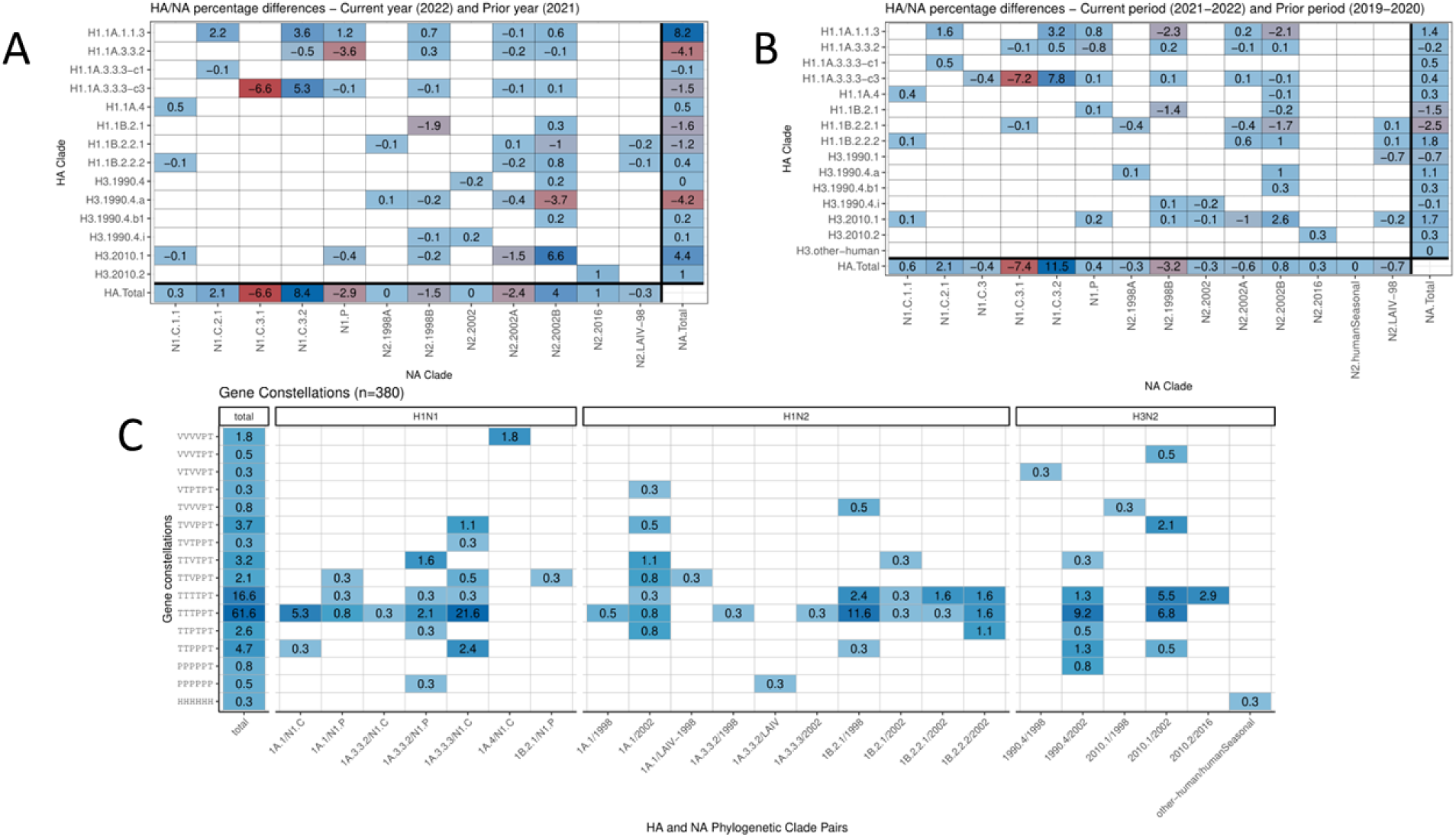
Representation of hemagglutinin (HA) and neuraminidase (NA) genetic clade pairings with whole-genome-sequence gene constellations collected within the United States Department of Agriculture influenza A virus (IAV) swine surveillance samples for 2022. The data are presented as percentages calculated based upon phylogenetic analysis and classification of segment sequences. The PB2, PB1, PA, NP, M, and NS genes were classified to evolutionary lineages of TRIG (T), Pandemic (P), or LAIV-related (L). The total percentage of each gene constellation observed within the 2022 data are listed as “total.”

### Genome constellation diversity and detection frequency

From 2009-2022, the three most common genome constellations were TTTPPT (32.1%, with 24.6% H1-1B.2.1/N2-1998B as the primary HA/NA pairing), TTTTPT (29.3%, with 14.2% H1-1B.2.2.1/N2-2002B as the primary HA/NA pairing), and TTPPPT (11.9%, with 64.5% H1-1A.3.3.3-c3/N1-C.3 as the primary HA/NA pairing). The remaining 26.5% of detections consisted of an additional 49 genome constellations that were paired with 133 different HA/NA combinations. From 2017-2022, the three most common genome constellations were TTTPPT (39.1%, with 28.4% H1-1B.2.1/N2-1998B as the primary HA/NA pairing), TTTTPT (30.0%, with 21.1% H3-2010.1/N2-2002B as the primary HA/NA pairing), and TTPPPT (13.9%, with 63.6% H1-1A.3.3.3-c3/N1-C.3 as the primary HA/NA pairing). The remaining 18.3% of detections consisted of an additional 36 genome constellations that were paired with 91 different HA/NA combinations. The three most common genome constellations detected during 2022 were TTTPPT (62.4%, with 32.1% H1-1B.2.1/N2-1998B as the primary HA/NA pairing), TTTTPT (14.5%, with 32.7% H3-2010.1/N2-2002B as the primary HA/NA pairing), and TVVPPT (11.4%, with 87.8% H3-2010.1/N2-2002B as the primary HA/NA pairing). The remaining 11.7% of detections in 2022 consisted of an additional 11 genome constellations that were paired with 29 different HA/NA combinations.

The time-series decomposition of weekly detections from 2009 and 2022 showed a general increase in IAV detections with a positive slope (Figure S1). Seasonal trends included an autumn IAV detection increase starting at approximately Week 40 (early October) peaking at Week 44 (early November), followed by a decline towards a low point at Week 52 (end of the year), a modest spring increase, and a decrease towards a minimum at Week 29 (mid-late July).

### Variable persistence and emergence of genome constellations in U.S. swine

The genome constellation persistence plot (Figure 3) revealed 297 unique combinations of genes, with 118 persisting for longer than one year, and 135 only being detected once. The average detection duration for a genome constellation was 1.8 years, although there were 8 combinations, e.g., H1-1A.3.3.2/N1-C.3PPPPPP and H1-1A.3.3.3-c3/N1-C.3TTTTPT, that persisted 10 years or more. Notably, ∼99% of M gene detections from 2017 to 2022 were from the H1N1pdm09 evolutionary lineage, and ∼80% of all data collected in 2022 had two or more genes related to the H1N1pdm09 evolutionary lineage. The detection of new genome constellations occurred at variable rates. Relatively few novel combinations were detected between the years of 2013 and 2017 (16.0 novel genome constellations/year). The periods before and after these years produced nearly two times as many new combinations per year (2010-2012, 28.7 novel genome constellations/year; 2018-2022, 28.0 novel genome constellations/year). The 2018-2022 period coincided with the introduction of a LAIV in the U.S. (28), which introduced new evolutionary lineages (V lineage designation) that were detected with varying persistence during this period. There were 14 states with positive source-sink scores that represented net generators of novel genome constellation diversity (Figure 4: CA, IL, IN, MN, MO, MT, NC, ND, OH, OK, PA, SD, TN, TX). AL and AZ had a source-sink score of 0. The remaining 10 states (CO, IA, KS, MI, NE, SC, UT, VA, WI, WY) had negative source/sink scores and reflect U.S. state locations where genome constellation diversity was detected for the final time, i.e., extinction of that genome constellation.

**Figure 3.**
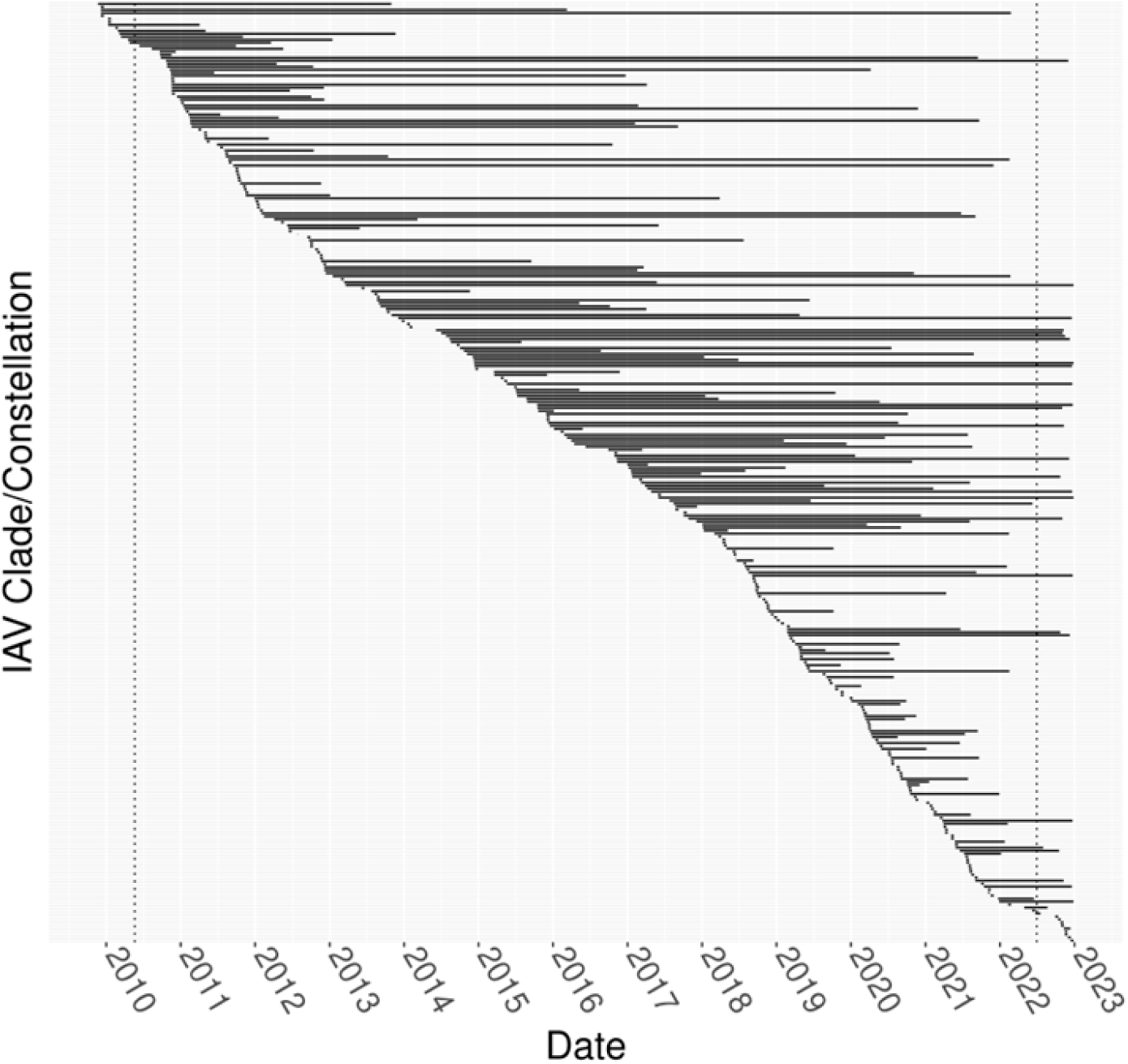
Persistence of influenza A virus genome diversity measured using hemagglutinin and neuraminidase genetic clade paired with evolutionary lineage for the remaining six genes. Persistence was denoted with a black bar connecting the first and last dates of the clade/constellation detection. Vertical dashed lines denote 180-day windows from the first and last detection dates in the dataset. Detections in the first and last 180 days of the dataset were not counted for source or sink event analysis. Viruses were ordered by date of first detection and a figure with clade/constellation in searchable form is provided in the supporting data.

**Figure 4.**
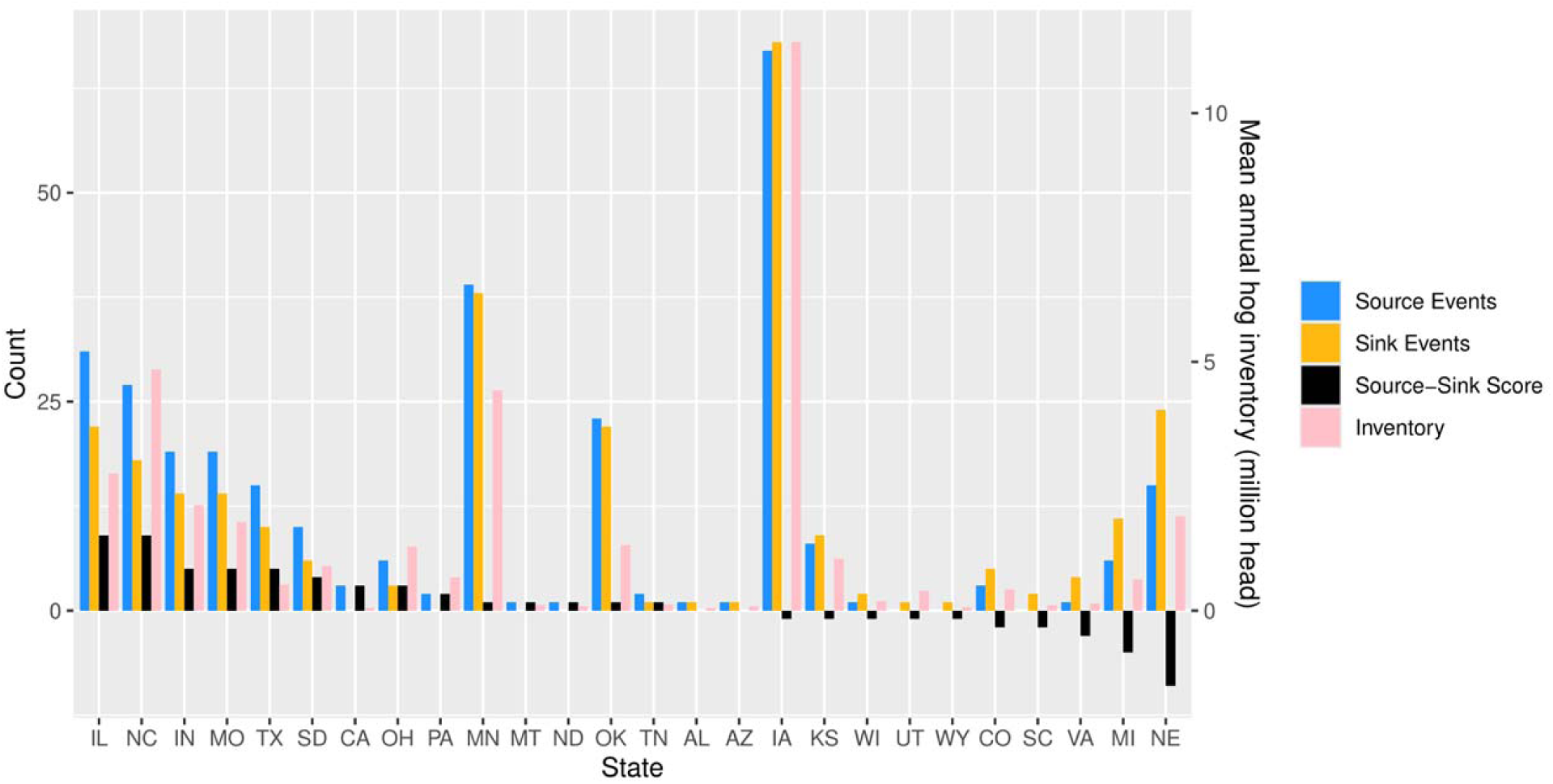
Counts of source and sink genome constellation detection events and mean swine inventory across swine-producing states for the period between 2010 and 2022. Scores were calculated for source/sink events of novel combinations of clades and gene constellations and the data in the graph were sorted by the source-sink score.

### Spatial and temporal patterns of genome constellation diversity

Zeta diversity was measured for each year between 2009 and 2022 and represented by the area under the line in Figure 5. In relative terms, Zeta diversity was low from 2009 to 2015, high from 2016 to 2020, and had a decreasing trend from 2020 to 2022. The ratio of Zeta diversity decline was best fit by the power law regression in years 2012, 2014, and 2022, suggesting that diversity patterns in these years were not driven by random processes, whereas all other years better fit the exponential regression. When all data were analyzed as a single group, the ratio of Zeta decline better fit the exponential regression (exponential regression AIC = -9.85, power law regression AIC = -6.76, Figure S2).

**Figure 5.**
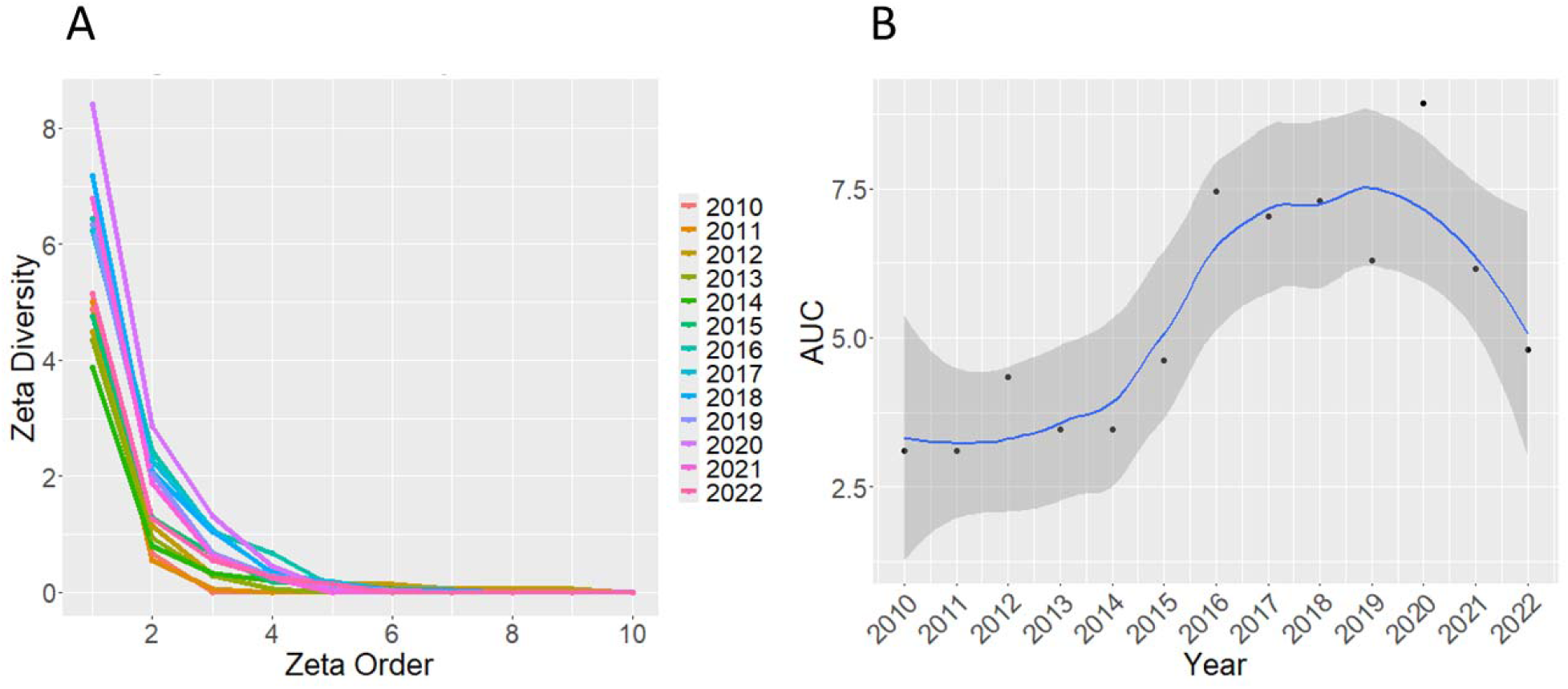
Zeta diversity per year for influenza A virus in swine between 2010 and 2022. High Zeta diversity at a given Zeta order means high proportion of the viruses were spread across a number of U.S. state locations equal to that Zeta order. High Zeta diversity at higher Zeta orders means that viruses are widely distributed. Low Zeta diversity at higher Zeta orders means that viruses are narrowly distributed. (A) Zeta diversity decreases most between orders 1 and 2 and approaches 0 by order 5. The shape of Zeta diversity decay changes little over time. (B) Area Under the Curve (AUC) of the Zeta curves in Figure A. Higher AUC denotes higher Zeta across all orders. AUC data are fit with a loess-smoothed line. Zeta diversity was low 2010-2015, high in 2016-2020, peaked in 2020, and started falling in 2021.

Markov chain models were applied to assess probabilities of pairwise interstate transmissions (Figure 6). For most U.S. states, following the detection of a virus, the subsequent detection was most likely to be within the same state, i.e., the diagonal value has the highest probability for that state. Specifically, the mean probability of the next detection being in the same state as the prior detection was 19.8%. The states with the highest probability of receiving new genome constellations were IA (20.9%), IL (11.1%), OH (9.73%), MN (9.30%), and IN (8.71%). The average probability of a given detection being the final detection before a sink event, i.e., the final state in the detection sequence being “XX”, was 9.52%.

**Figure 6.**
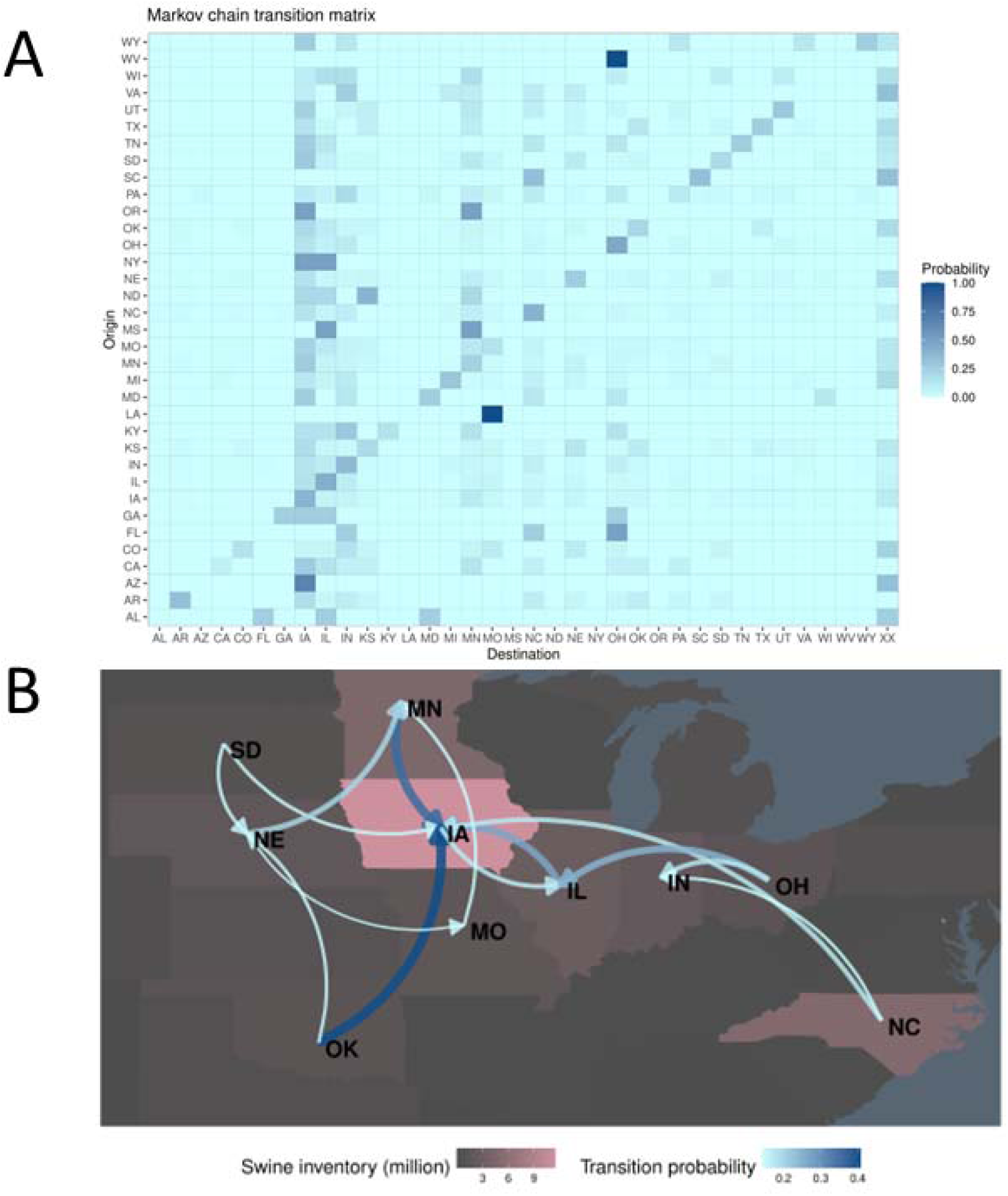
Markov chain model trained on state detection order of influenza A virus (IAV) clades and genome constellations. State detection order was used as an estimate of IAV interstate dispersal. (A) Transmission matrix heatmap describing the probability that a novel genome constellation will disseminate or stay within the location. The full model permitted repeated serial detections within a given state, and the probability values quantify whether a novel constellation/clade combination would move from an origin (rows) to a destination (columns). The “XX” represents the condition in which the constellation/clade combination terminated - a sink event. (B) A network graph reduced representation of a Markov chain model not permitting serial detection within states. This graph focuses on the 10 states with the highest swine inventory, plotted with background color, and only includes edges with a value of 0.15 or greater, with transition probability indicated with line width and color.

The power of this model was evaluated based on its performance in predicting outcomes in given scenarios. The model predicted the final state a strain would spread to with 20.6% accuracy, or 34.2% when state sequences were reduced to the top 10 swine-producing states. The model predicted the state of final detection with 37.7% accuracy, or 44.6% when state sequences were reduced to the top 10 swine-producing states. We also applied this model to predict extinction events and success/failure to spread to new states and contrasted these results against the strategies of always guessing further detection or spread (“assume not XX”) or always guessing extinction or failure to spread (“assume XX”). This model predicted extinction/persistence with 54.7% accuracy for strains with a single prior detection and 76.6% accuracy for strains with two prior detections. In contrast, the “assume not XX” strategy had prediction accuracies of 57.0% for strains with a single prior detection and 76.6% for strains with two prior detections. This model predicted interstate spread success/failure with 55.1% for strains previously detected in a single state and 48.6% accuracy for strains previously detected in two states. In contrast, the “assume XX” strategy had prediction accuracies of 57.0% in the first comparison, and the “assume not XX” strategy had a prediction accuracy 65.9% for the second comparison.

## DISCUSSION

Influenza A virus infection in swine is a recurring problem with year-round circulation and seasonal epidemic peaks in October-November and March-April. The cocirculation of multiple distinct genetic clades with frequent reassortment makes controlling IAV via vaccination difficult. A multivalent vaccine to reflect diversity within a single region may require four to six H1 and two to three H3 components to cover observed diversity (40). These vaccine components risk becoming less relevant given the constant transport of pigs and their IAV across the U.S. (23, 41–44). Further, the introduction of unique HA/NA and genome constellations to new locations may trigger evolution via reassortment, with associated mutations in the surface glycoproteins that affect antigenic phenotype and reduce vaccine efficacy (41). Consequently, controlling diversity in locations that appear to seed diversity to other regions would be beneficial. The inclusion of vaccine antigens that are less frequently detected in genomic surveillance may have the advantage of preemptively removing novel genetic clades and genome constellations prior to dissemination. This could reduce HA/NA genetic diversity, reduce the impact of reassortment on virus phenotype, and enhance control programs that seek to match antigenic components in vaccines to circulating diversity.

The USDA National Surveillance Plan for Swine Influenza Virus in Pigs has provided passive monitoring of the genetic diversity and evolution of IAV in U.S. swine for nearly 15 years. These data have been used to demonstrate a critical interface between people and pigs, with reverse zoonoses in the 1990s, 2010s, and 2020s establishing four H3 lineages (8, 45) and near-constant human-to-swine H1N1pdm09 virus introductions affecting genome constellation patterns (46). In the past two years, there have been significant shifts in observed genetic diversity (Figure 2). Of the 14 HA clades that were detected in 2022, only three HA clades increased in detection frequency from the prior year, five decreased, and five had trivial changes in detection frequency. Of the three increasing clades, the 1A.1.1.3 HA clade was associated with a reassortment in the neuraminidase (NA) gene. Prior to 2020, the 1A.1.1.3 clade was paired with the N2-1998B, but from 2021 onwards it was most often paired with N1-Classical (N1-C.1.1 or N1-C.3.2) or N1.PDM NA genes. Similarly, in 2022 there was a significant increase in the detection frequency of the H3-2010.1 lineage, with the concurrent decrease in detection frequency of the H3-1990.4.a, but these HA lineages shared a common NA gene and internal gene constellation. These data suggest that reassortment may result in novel gene pairings that affect transmission phenotype (31).

Variation in host immunity to different HA genes may allow for the persistence of rare clades in different populations that may then increase in frequency as the rare viruses are introduced into naïve hosts through animal movement (30). This proposition follows an established metacommunity theory framework whereby local and regional processes, including dispersal, operate to mediate diversity patterns (47). Currently, there is one commercially licensed vaccine product for swine IAV (11), and many swine production systems rely upon custom or autogenous vaccines derived from genomic surveillance. Our analysis, as well as prior research (17, 42), indicates relatively frequent movement of pigs and their IAV to new locations. The lag in time between identifying HA diversity within a swine herd and the production of a matched custom or autogenous vaccine likely creates a variable immunological landscape that allows time windows in which HA clades may be more or less successful in the swine population (30). It is plausible that the fluxes we report in HA clade detections are associated with a heterogenous immune landscape determined by changes in vaccine-driven or natural immunity.

Our data reveal shifts in evolutionary dynamics over time. In 2021 there were 32 novel genome constellations detected, the most unique combinations in a single year since 2011, in which there were 33. There was relatively limited genome constellation novelty from 2013 to 2017. In 2012-13, Porcine Epidemic Diarrhea Virus (PEDV) emerged and caused an epidemic in the U.S. swine herd (48). There were no adequate vaccine strategies for PEDV in the initial epidemic, and the swine industry responded with stringent biosecurity practices (49). An indirect consequence of PEDV control may have been a reduction in pig movement and fewer opportunities for IAV reassortment. Immediately following this period, there was an increase in new genome constellations driven by the use of and reassortment with a live attenuated influenza virus vaccine (28). Conversely, periods in which no novel combinations were detected may reflect changes in methods in the USDA’s IAV surveillance plan. More broadly, these time windows illustrate that evolutionary rates are not stable over time, and that (re)introduction of novel or previously absent genetic diversity and subsequent reassortment can increase the observed rate of IAV diversification and may be associated with IAV-nonspecific biosecurity practices (50).

We find that most new IAV genomic diversity is first detected in Midwest states (IL, MO, IA, among others), and we refer to these states as net sources of swine IAV diversity. Conversely, states with negative source-sink scores are net sinks of diversity. While the number of sink events and source events were positively correlated with swine inventory per state, the sign of the source-sink score was not explained by pig population. Rather, a state’s role as a source or a sink of diversity is likely tied to its location in a network of swine transport. To quantify this, we used Markov chain models to assess the geographic spread of novel IAV following its initial detection. The resulting transition matrix and network graph provided a probabilistic framework suggesting that the location where a novel genome constellation is first detected impacts its persistence and spread. Because the USDA IAV swine surveillance plan has targeted cases based upon clinical compatibility, there is no information on negative cases, and it does not collect fine-scale information on pig movement and the abiotic environment available that could help to refine this modeling approach. Future statistical forecasting efforts must overcome these challenges to effectively and reliably predict whether newly reassorted IAV detected in swine will disperse across the U.S. Further, while the links in our dispersal network likely approximate the ‘swine-ways’ across which pigs and their IAV are transported across the United States (42), a model trained on interstate swine transport data that is nested within a phylodynamic framework will empower more accurate predictions of IAV dissemination.

We also documented variation in diversity patterns across each year of our study. Zeta diversity at higher orders (2–6) was low from 2010-2016, then Zeta doubled for several years, and then began to fall again. In some years, the distribution of IAV diversity in swine was random; while in other years, it was so narrowly distributed that it presented a statistical signal associated with local adaptation, i.e., each location had unique IAV in swine, e.g., in 2012-2014, IAV was narrowly distributed. Local adaptation in this scenario likely reflects host immune status, as many producers use custom vaccines that reflect the IAV diversity in their production system. This targeted vaccination may restrict which IAV is detected and sequenced within the farm and within a region, reduce co-infection and reassortment, and restrict the emergence of new genome constellations.

Our analyses also identify regions where surveillance may be more targeted, as they act as epicenters for reassortment of IAV in swine. Surveillance in these regions may be able to identify novel reassorted IAVs that impact animal health and have increased zoonotic potential. This suggestion is consistent with the prior work that argued that vaccine composition should consider IAV diversity within states and swine production regions in addition to aggregated national patterns (3). More generally, using reassortment frequency as a supplement to HA/NA phylogenetic clade identification for vaccine control efforts would be beneficial, as there is some evidence that IAV surface proteins evolve more rapidly following reassortment (17). Additionally, new detections of genome constellations can be flagged for further biological characterization to identify whether specific genes contribute to transmission and/or pathology. These data also identify which segment combinations persist, offering insights into coevolution and its effects on virus transmission and pathology.

Our results show that different states play disparate roles in the evolution of IAV in swine in the U.S. Some states serve as hotspots of IAV reassortment and contribute significantly to the spread of novel variation to the rest of the U.S., while others seem to serve as major transit hubs in the dispersal of IAV from one state to another. More nuanced models tracking the flow of IAV across the U.S. may aid in directing control measures to regions where they would be most impactful. Our results also show that the rate of diversification in IAV is not consistent across time. The 2013-2017 period showed a marked reduction in novel reassortant diversity. While the factors influencing changes in diversification rate are yet to be discovered, the fact that this rate is not fixed means that human intervention may be able to lower this rate, to the benefit of the health of swine and the public. Consequently, our data demonstrate that a more nuanced view to tracking the evolutionary patterns of IAV may allow us to move toward control methods that reduce observed genetic diversity in the surface proteins and may facilitate our understanding and ability to predict host responses to infection.

## ACKNOWLEDGMENTS

We gratefully acknowledge pork producers, swine veterinarians, and laboratories for participating in the USDA National Surveillance Plan for Swine Influenza Virus in Pigs and publicly sharing sequences. This study was funded by: USDA-APHIS (ARS project number 5030-32000-120-80-I); USDA-ARS (ARS project number 5030-32000-231-000-D); the National Institute of Allergy and Infectious Diseases, National Institutes of Health, Department of Health and Human Services (contract number 75N93021C00015); the USDA-ARS Research Participation Program of the Oak Ridge Institute for Science and Education (ORISE) through an interagency agreement with the US Department of Energy (DOE) (contract number DE-SC0014664); and the SCINet project and the AI Center of Excellence of the USDA-ARS (ARS project numbers 0201-88888-003-000D and 0201-88888-002-000D). The funders had no role in study design, data collection and interpretation, or the decision to submit the work for publication. Mention of trade names or commercial products in this article is solely for the purpose of providing specific information and does not imply recommendation or endorsement. USDA is an equal opportunity provider and employer.

## Data availability

All genetic sequence data are available at NCBI Genbank with USDA IAV in swine surveillance data within a searchable interface at https://flu-crew.org. The code associated with analysis in this manuscript is provided at https://github.com/flu-crew/iav-sources-sinks.

