## Supplemental Figures for "Sources and sinks of influenza A virus genomic diversity in swine from 2009 to 2022 in the United States"

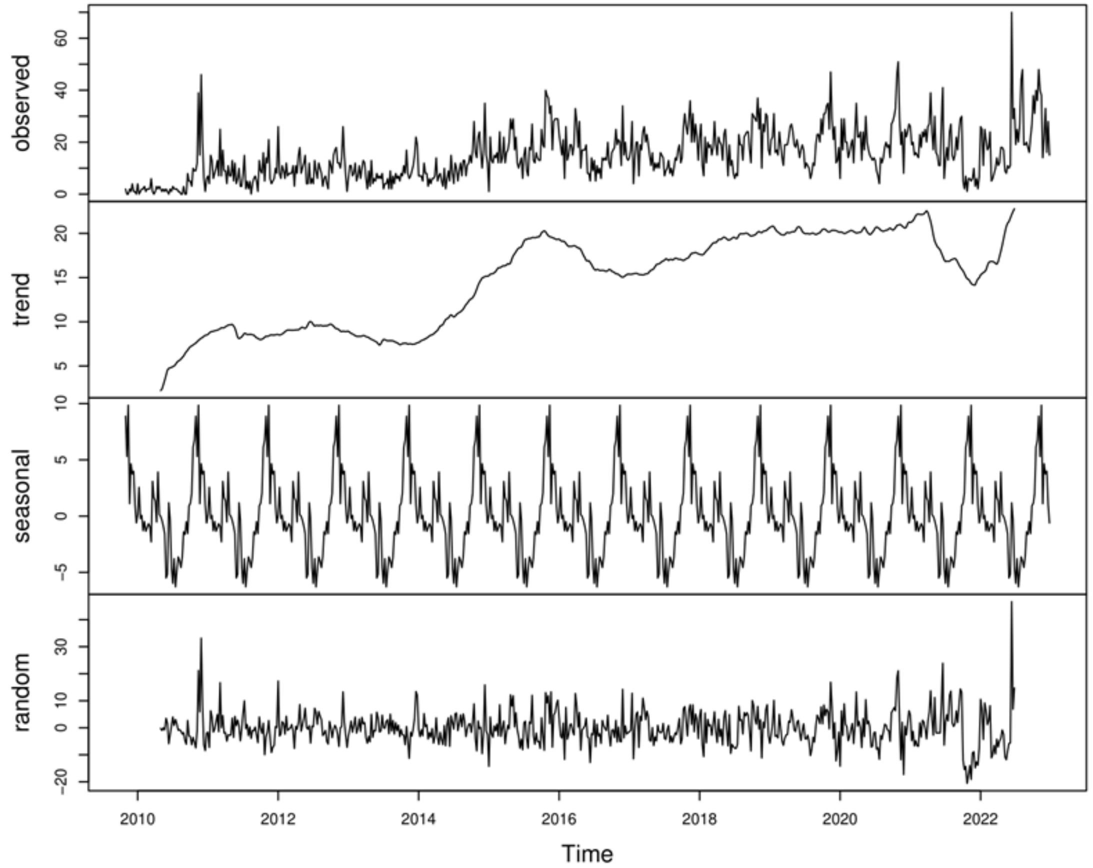


**Figure S1**. Temporal decomposition of detection counts (cases per week) of influenza A in swine. From the observed case counts, the moving average finds the overall trend, and the additive seasonal effect is found by averaging each week over all years. The trend and seasonal components are removed from the observed counts to calculate the remaining random noise. Over the data series, detections per week generally increase until 2020, and the average magnitude of seasonal effects is similar to that of random effects.

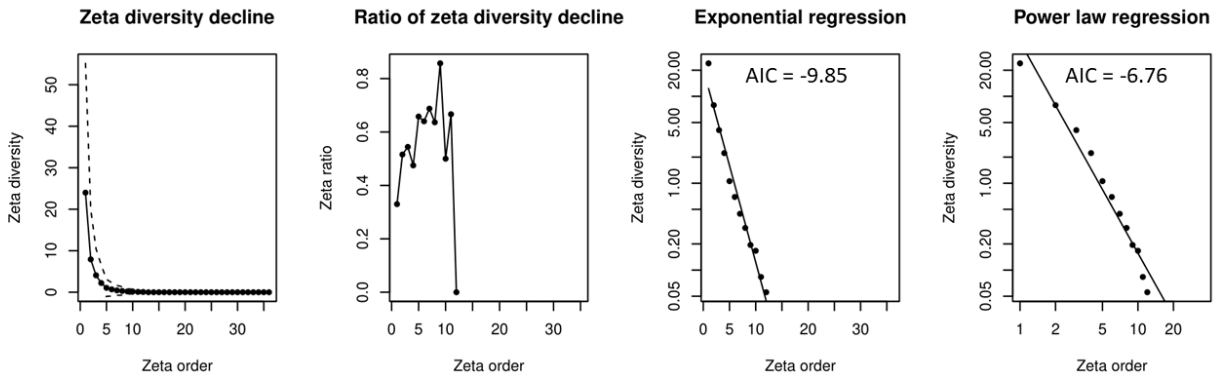


**Figure S2**. Zeta diversity from data from all years combined. The first panel shows Zeta diversity decay with increasing order; the second shows the ratio of retained diversity to lost diversity with each increase in Zeta order; and the third and fourth panels demonstrated the exponential and power law regressions of the ratio of Zeta diversity decline. AIC showed that the exponential regression fit the data more closely than the power law regression, indicating that Zeta diversity was not more narrow than null expectations, which was evidence against niche separation between states.
